## Supplementary figures and images for "Crowd-sourced observations of a polyphagous moth reveal evidence of allochronic speciation varying along a latitudinal gradient"

### Supplemental Figure 1

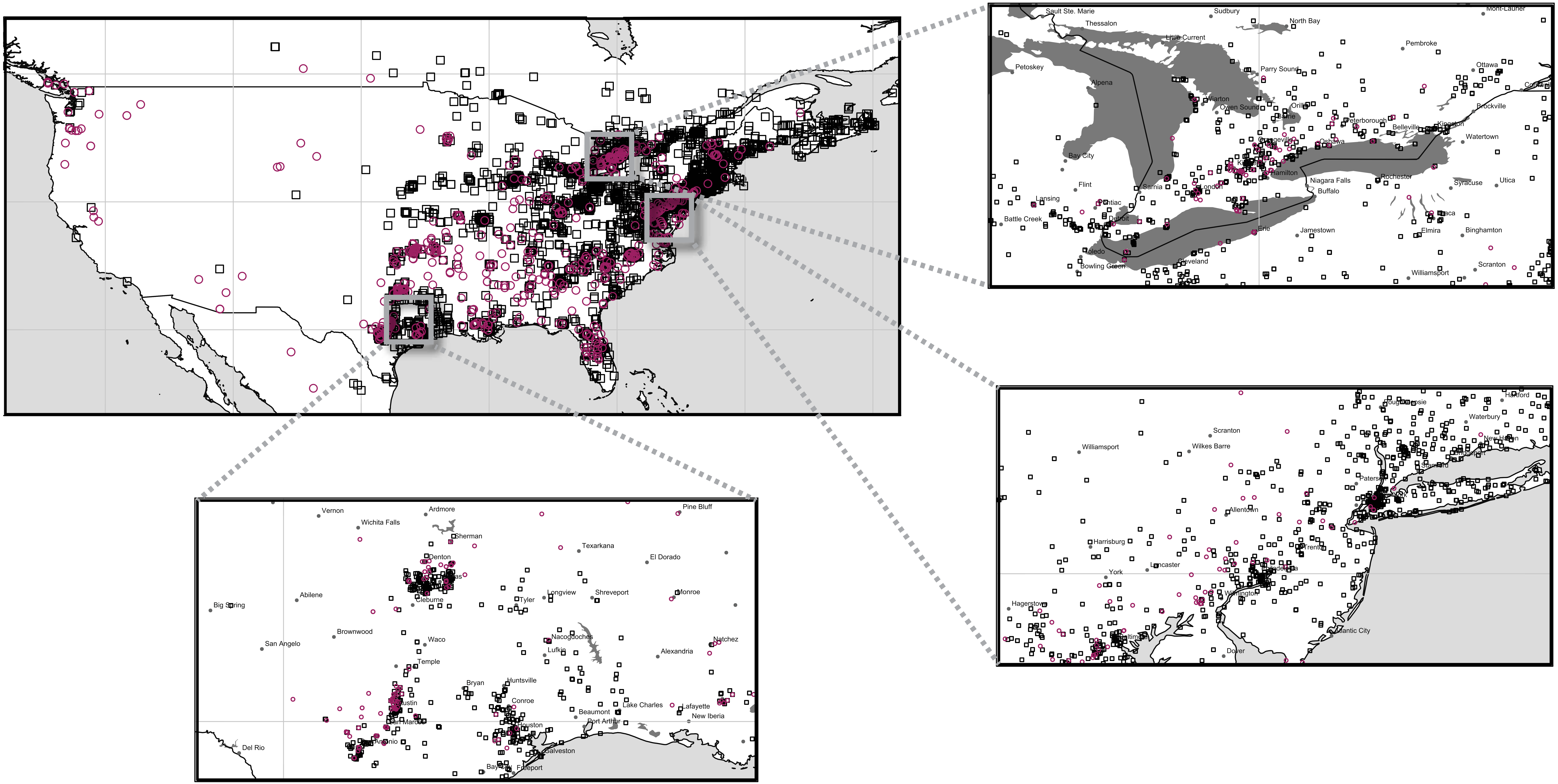

### Supplemental Figure 2

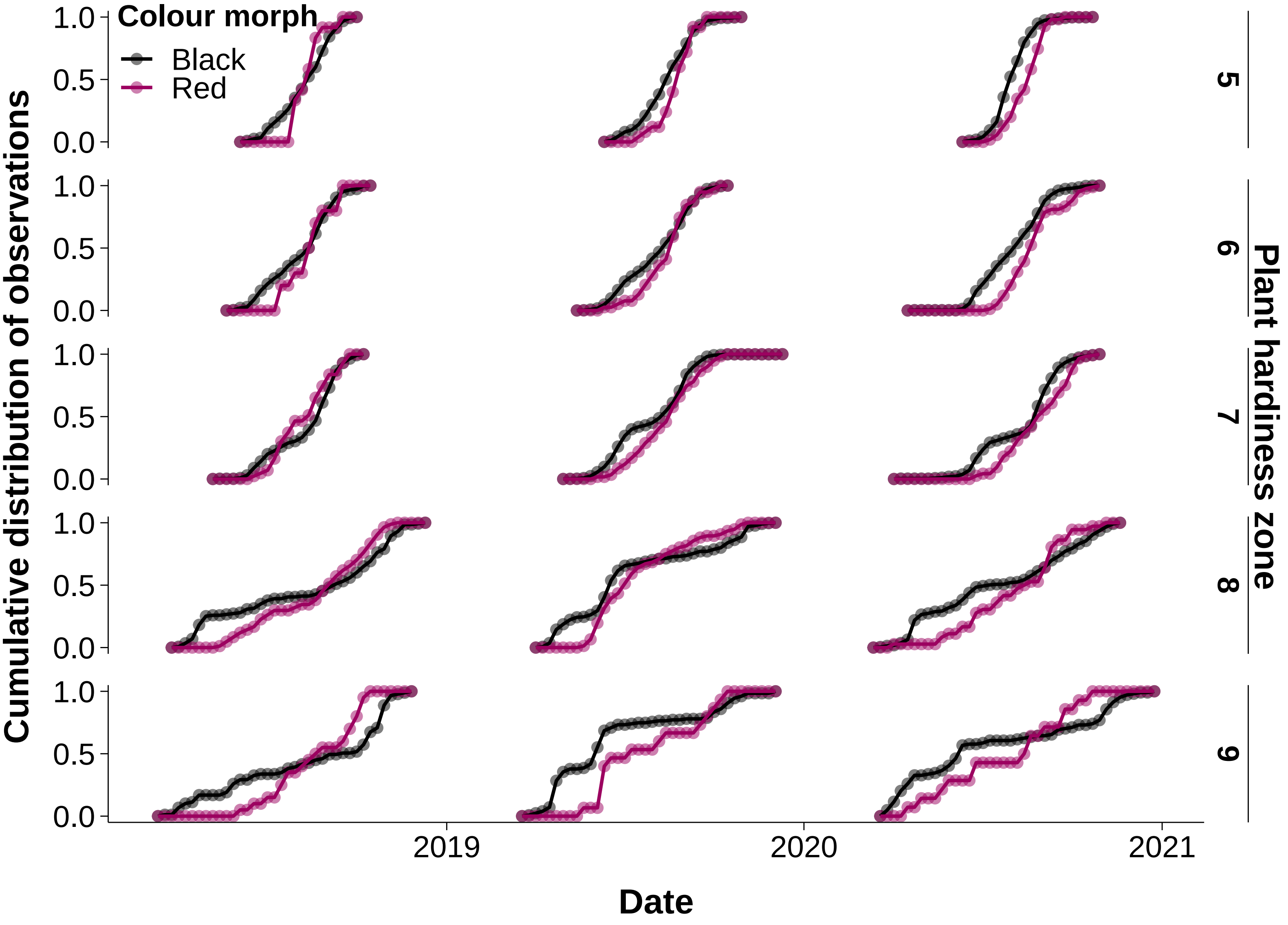

### Supplemental Figure 3

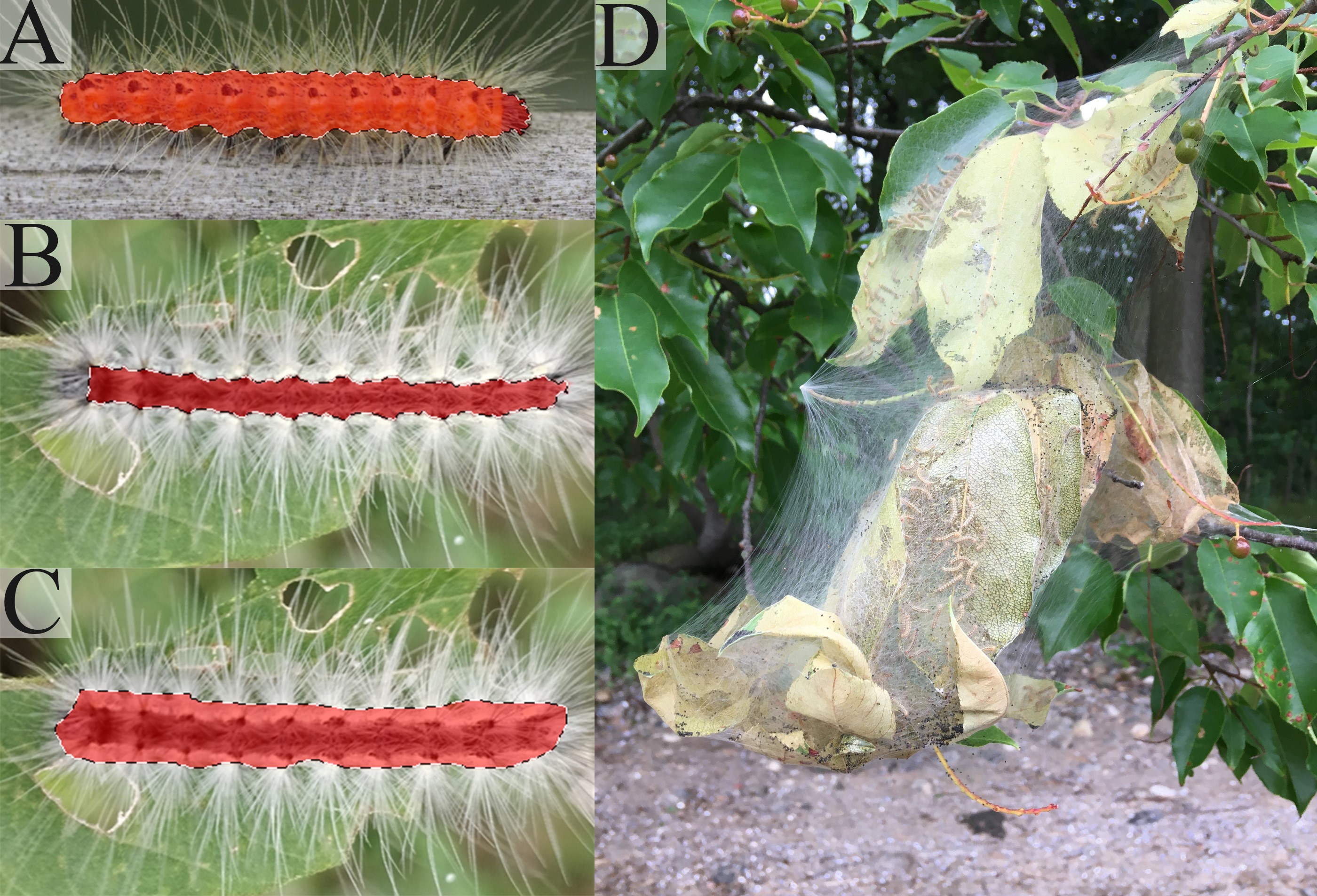

### Supplemental Figure 4

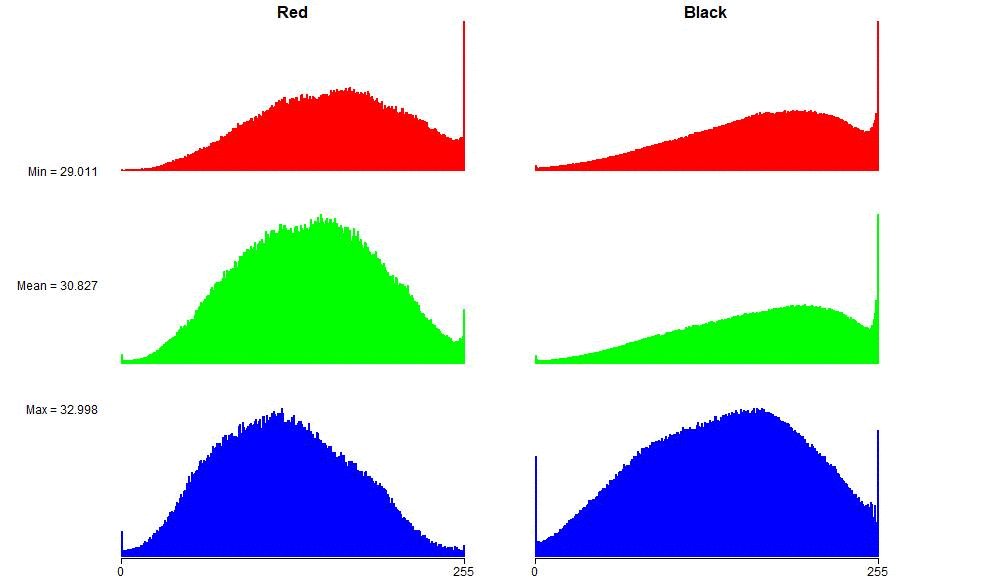
