## Supplemental image captions for "Crowd-sourced observations of a polyphagous moth reveal evidence of allochronic speciation varying along a latitudinal gradient"

**Supplementary Results**

Full-sized figures and formatted tables available at <https://osf.io/x64gp/?view_only=cb630bc7e24d487a9fdaa1a5682e82cb>

**Legend:**

**Table S1**. Coefficients of linear discriminants from LDA analysis of variables separating colouration of red and black morphs in iNaturalist photographs.

**Figure S1.** Regional distribution of iNaturalist fall webworm observations from 2018-2020 with colour morph indicated.

**Figure S2.** Cumulative distribution functions of red and black observations in each year and plant hardiness zone.

**Figure S3.** Fall webworm image isolation and selection in Adobe Photoshop for colour phenotype analysis.

**Figure S4.** Animated RGB histogram of red and black fall webworm larvae across their latitudinal range.

**Table S1**. Coefficients of linear discriminants from LDA analysis of variables separating colouration of red and black morphs in iNaturalist photographs. Among the coefficients of linear discriminants, the mean green value, mean red value, and red standard deviation have the highest absolute values (-2.60, 1.75, and -0.97 respectively).

| *Variable* | *Value* |
| --- | --- |
| *GreenMean* | *-2.60732* |
| *RedMean* | *1.750011* |
| *RedSD* | *-0.96515* |
| *BlueMax* | *0.796165* |
| *RedMin* | *0.7282* |
| *GreenMax* | *-0.71824* |
| *BlueSD* | *0.656008* |
| *BlueSkew* | *-0.65438* |
| *RedSkew* | *0.638823* |
| *GreenKurt* | *0.512545* |
| *RedKurt* | *-0.3341* |
| *GreenMin* | *-0.3192* |
| *RedMax* | *0.300597* |
| *GreenSkew* | *0.163374* |
| *BlueMin* | *-0.112* |
| *BlueKurt* | *-0.1039* |
| *GreenSD* | *0.089111* |
| *BlueMean* | *0.077523* |

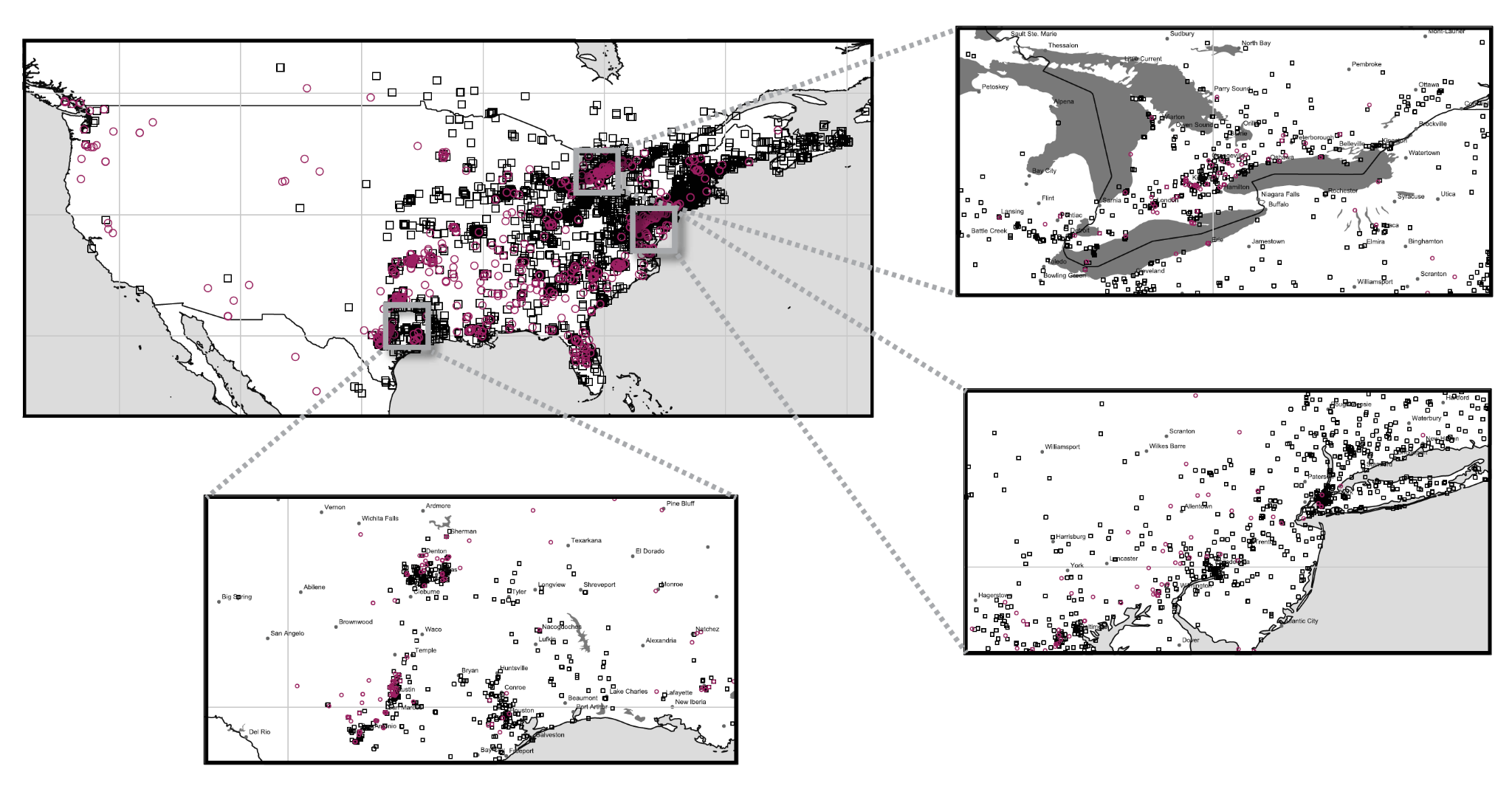

**Figure S1.** Regional distribution of iNaturalist fall webworm observations from 2018-2020 with colour morph indicated. Red and black morph fall webworms occur sympatrically at a regional and local level, implicating a sympatric form of speciation. Regions shown are southern Ontario, northeastern USA, and southern Texas.

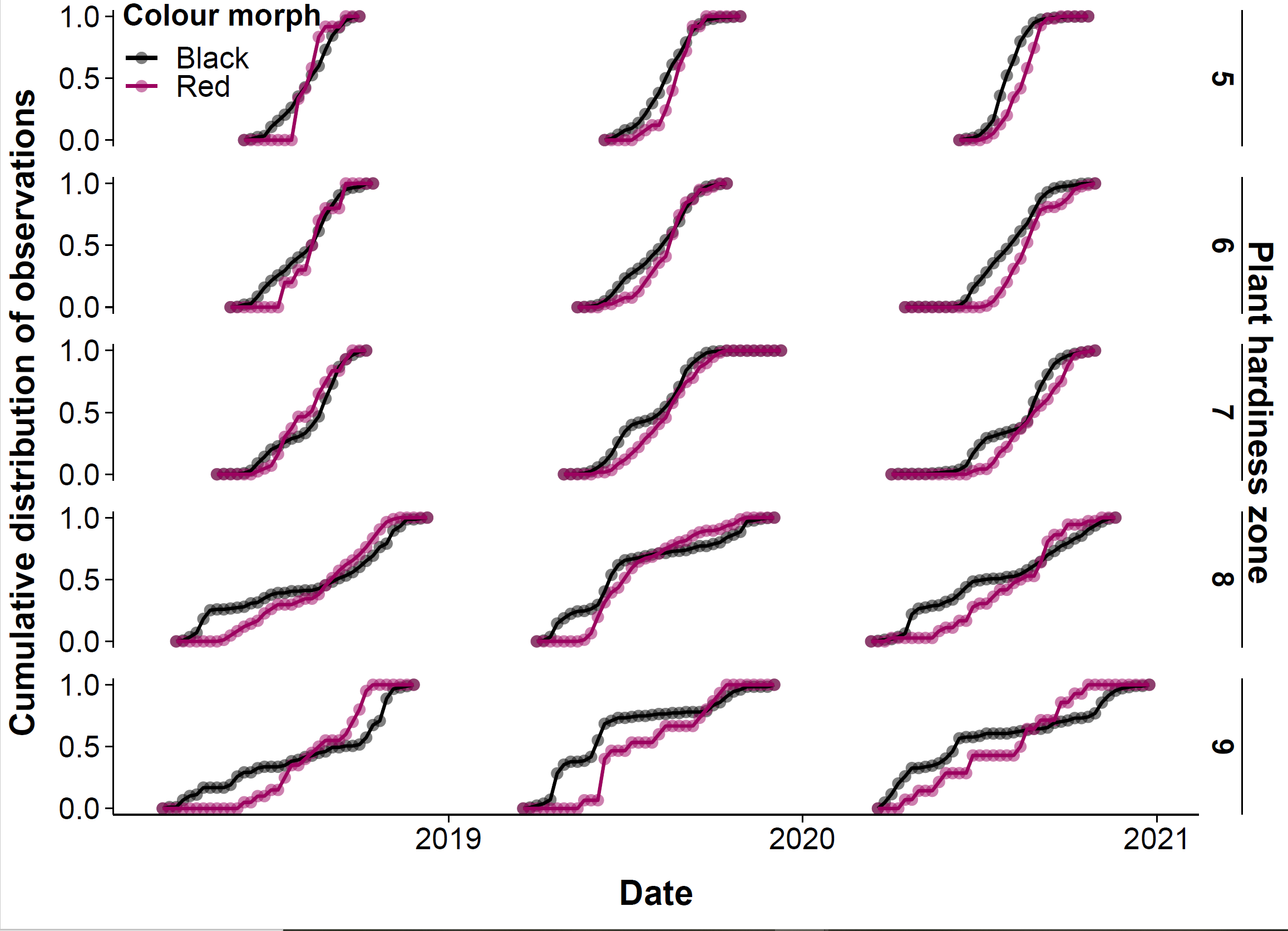

**Figure S2.** Cumulative distribution functions of red and black observations in each year and plant hardiness zone. Points indicate cumulative proportion of observations, while lines show trends in distribution across years.

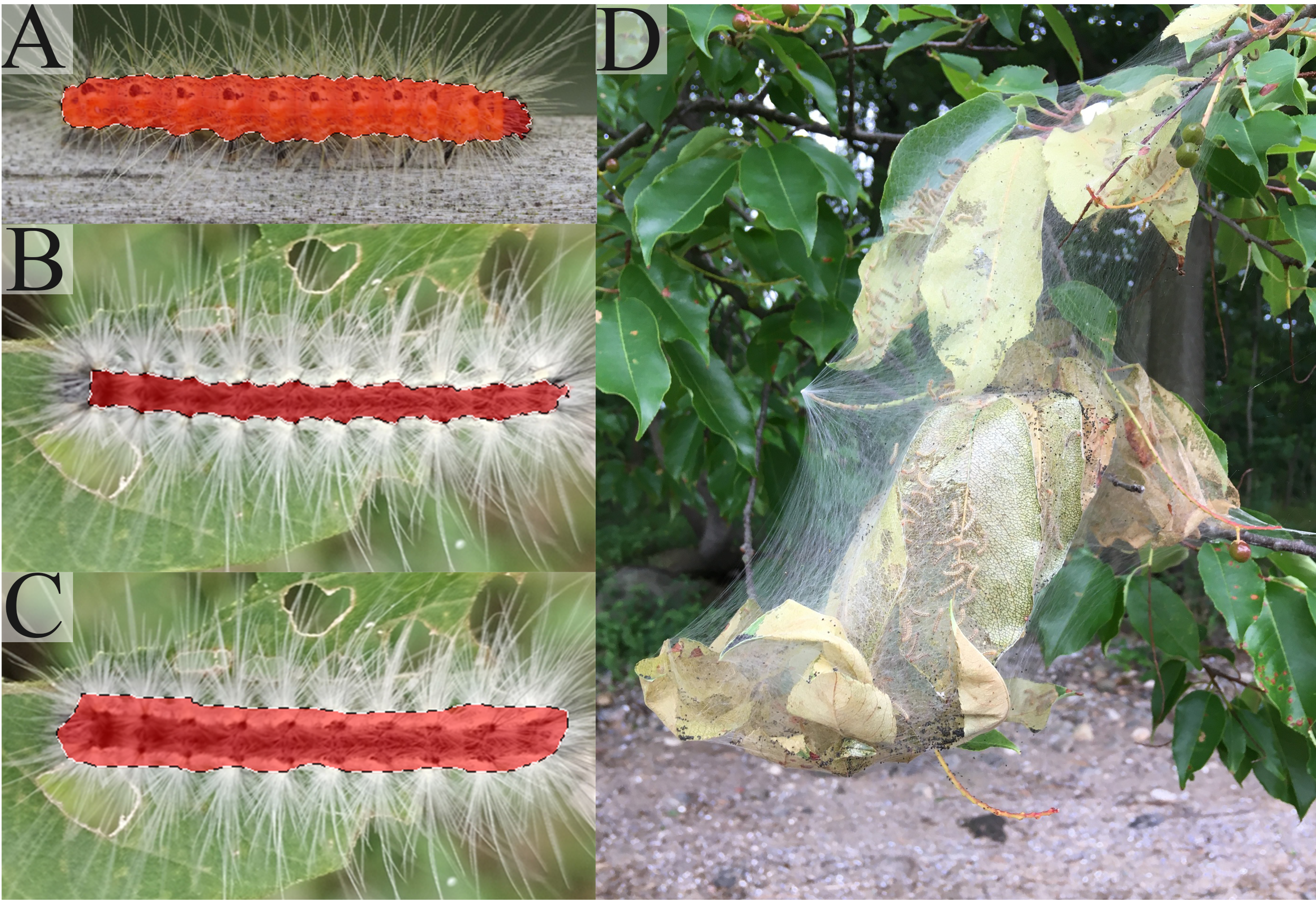

**Figure S3.** Fall webworm image isolation and selection in Adobe Photoshop for colour phenotype analysis. A) Complete isolation of fall webworm larva by Photoshop using object detection tool, no manual adjustment needed. B) Incomplete selection of fall webworm larva by object detection tool. Corrected manually by researcher using freehand lasso tool. C) Complete selection of fall webworm larva in Photoshop after manual correction using freehand lasso tool. D) Example of photo removed from colour phenotype analysis. Photo was low resolution, had poor exposure, was taken from a far distance, and had an object obstructing the webworms. Photos shown here obtained from iNaturalist.org under public domain licensing (CC0).

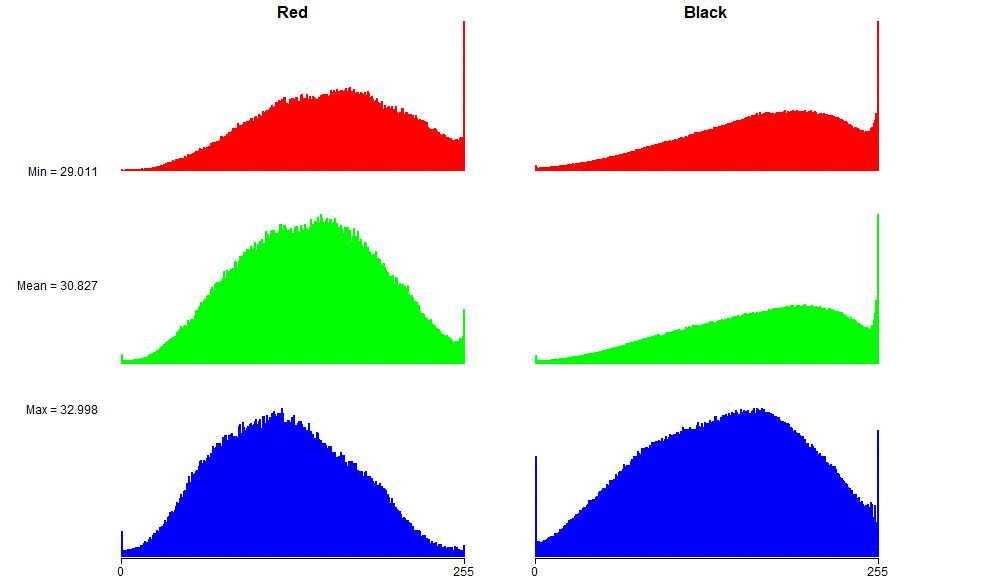

**Figure S4.** Animated RGB histogram of red and black fall webworm larvae across their latitudinal range.
